# A novel multicellular model of the adult mouse sinoatrial node retains spontaneous electrical activity and enables live investigation of the S100B-associated cell population

**DOI:** 10.64898/2026.07.17.735408

**Authors:** Georgiana Luisa Baca, Robert E. Monticone, Bruce D. Ziman, Shah Md Toufiqur Rahman, Pathik Parekh, Sadia Afrin, Christopher Dunn, Richard Telljohann, Dongmei Yang, Kwan-Wood Gabriel Lam, Peter Killeen, Dimitrios Tsitsipatis, Ana-Maria Zagrean, Myong-Hee Sung, Nigel H. Greig, Supriyo De, Allison B. Herman, Payel Sen, Rafael de Cabo, Edward G. Lakatta

## Abstract

Approximately half of the adult sinoatrial node (SAN) consists of non-myocyte populations, indicating that cardiac pacemaking depends on interactions within a multicellular tissue rather than on pacemaker cardiomyocytes alone. Among these, an S100B-associated cell population has been implicated in pacemaker function, yet its identity and physiological roles remain poorly understood. These cells are rare and dispersed throughout the small, structurally complex SAN, making them difficult to observe repeatedly while preserving the native multicellular environment. Here, we established a dissociated multicellular culture of adult mouse SAN tissue on soft collagen–gelatin hydrogels that retains spontaneous electrical activity and permits longitudinal live imaging of S100B-associated cells. Using an S100B-EGFP^+^ reporter, we identified at least six reproducible morphological and behavioral phenotypes, including migration, proliferation, phagocytic behavior, and spontaneous self-organization into three-dimensional clusters. Cultures remained spontaneously electrically active for more than 10 days in vitro, with peak activity around day 10. This multicellular culture model bridges the gap between intact SAN preparations and isolated-cell cultures, allowing repeated observation of rare S100B-associated cells within a spontaneously active multicellular environment.

**Highlights:**

- The platform enables longitudinal live imaging of rare S100B-associated cells within an diverse multicellular SAN culture.
- Live imaging reveals at least six reproducible morphological and behavioral phenotypes of S100B-associated cells.
- Dissociated multicellular SAN cultures remain spontaneously electrically active for more than 10 days in vitro.

## Introduction

Cardiac pacemaking emerges from interactions among multiple cell types within the sinoatrial node (SAN), yet the identity and behavior of endogenous S100B-associated cells remain poorly understood.In the adult human SAN, the heart’s primary pacemaker, HCN4^+^ pacemaker cardiomyocytes generate rhythm through coordinated membrane ion currents and intracellular Ca^2+^ cycling [1], yet nearly half of the tissue volume consists of non-myocyte populations [2]. Single-nucleus and spatial transcriptomic studies reveal a structured microenvironment comprising fibroblasts, macrophages, endothelial and adipocyte populations, autonomic neurons, and glial-associated interstitial cells [3,4], organized into distinct anatomical domains with a central pacemaker core surrounded by specialized stromal and immune compartments [4].

These non-myocyte populations actively contribute to pacemaker function. Fibroblasts form coupled networks with pacemaker cardiomyocytes, regulate diastolic depolarization and beat-to-beat interval [5,6], and provide contact-dependent signals required for normal pacemaker metabolism and electrical activity [7]. The extracellular matrix composition and tissue mechanics further influence SAN automaticity [8,9], while macrophages, neurons, and glial-associated cells contribute to nodal structure and function [10,12,13].

Among these non-myocyte populations, S100B is of particular interest because it has been implicated in SAN function. Three-dimensional imaging identified a peripheral glial network and an S100B^+^ interstitial population closely associated with the HCN4^+^ pacemaker meshwork [14– 17], while exogenous S100B disrupts pacemaker Ca^2+^ synchrony and increases rhythm variability [14]. Beyond the heart, S100B functions as an activity-dependent glial signal that modulates neural network activity, and cardiac glia release S100B following injury [18–23].

The identity, organization, and physiological roles of endogenous S100B-expressing cells remain poorly defined in the heart. S100B is not cell-type specific, and previous single-cell transcriptomic studies did not identify S100B as a specific marker of any discrete SAN cell population [24,25]. Studying these cells requires experimental models that preserve the native multicellular environment while allowing longitudinal observation of this rare, dispersed cell population.

The SAN is small and structurally complex, making repeated observation of rare dispersed cell populations while preserving their native multicellular environment technically challenging. Existing in vitro models either isolate individual cell populations [26] or generate pacemaker-like cells from stem cells, sacrificing the native multicellular context, a limitation for studying how the node’s diverse cell types interact. Here, we developed a dissociated multicellular culture of the adult mouse SAN using an S100B-EGFP^+^ reporter, live-cell imaging, and network electrophysiology

## Materials and Methods

### Animals

Young wild-type C57BL/6J mice (JAX #000664) were used initially to optimize imaging platforms and cell plating conditions, and as non-reporter controls. Male and female B6;D2-Tg(S100B-EGFP^+^)1Wjt/J mice (JAX #005621; the “Kosmos” S100B-EGFP^+^ reporter line), which express EGFP^+^ under the human S100B promoter, were used at 12–24 weeks of age. This strain enables direct visualization and live tracking of S100B-expressing cells without additional labeling. All procedures were approved by the Animal Care and Use Committee of the National Institute on Aging, NIH (protocol 457-LCS-2027). Mice were anesthetized with sodium pentobarbital, hearts were rapidly excised, and the SAN was isolated under light microscopic guidance.

### Enzymatic dissociation of SAN tissue

Freshly dissected SAN tissue was enzymatically dissociated to obtain single-cell suspensions suitable for culture and downstream analysis. For each preparation, SAN tissue from two B6;D2-Tg(S100B-EGFP^+^)1Wjt/J mice were pooled and subjected to collagenase-based digestion optimized for small, low-yield cardiac regions, yielding reliable, viable suspensions containing both S100B^+^ and S100B^−^ cell populations.

### Hydrogel-based culture platform

SAN-derived cultures are cellularly heterogeneous, so we matched substrate to purpose, balancing biomechanical physiology with optical performance. We benchmarked rigid polystyrene, glass-bottom dishes, and collagen and gelatin hydrogels (Softview; Matrigen, SF-35-20). Hydrogels were prioritized for tissue-like compliance: polystyrene is non-physiological (~2 GPa), whereas the SAN matrix is ~17 kPa and collagen and gelatin hydrogels span the physiological range (~1–14 kPa). Compliant substrates favor automaticity (increased Hcn4, reduced Cx43, slower Ca^2+^-wave propagation) and keep cardiac fibroblasts quiescent, whereas stiff plastic drives myofibroblast conversion [27,28,29]. We used Softview plates (~6–14 kPa) for live and time-lapse imaging and glass-bottom MatTek and polimer iBIDI dishes for high-resolution immunofluorescence. Coating conditions and seeding densities were benchmarked across substrates; the morphological heterogeneity reported here was reproducible across substrates.

### Live-cell calcium imaging

SAN-derived cells were cultured on compliant hydrogel and collagen substrates and loaded with a membrane-permeable calcium indicator (Calbryte 520 AM). High-resolution time-lapse imaging of intracellular Ca^2+^ dynamics was performed on a Zeiss AxioExaminer D1 upright microscope equipped with a sensitive sCMOS camera and controlled excitation and emission filters. Imaging conditions minimized phototoxicity, and temperature and gas environment were tightly regulated.

### Time-lapse and live nuclear imaging

Long-term monitoring (24–72 h) of SAN cultures was performed using Keyence BZ-X810 and Zeiss LSM 880 microscopes with environmental control. Collagen-coated hydrogel and polymer-bottom chambers were used to assess spontaneous cellular activity, cluster formation, and migration. Imaging intervals ranged from 5 to 15 minutes, with settings optimized to minimize photobleaching. For live nuclear labeling during imaging, cultures were incubated with the cell-permeable, non-toxic far-red DNA probe SPY650-DNA (Spirochrome; excitation/emission 652/674 nm), applied at 1:1000 from a 1000× DMSO stock without a wash step. SPY650-DNA is pseudocolored red in fluorescence overlays and is distinct from the NucBlue counterstain used on fixed preparations.

### Immunofluorescence profiling

Primary SAN cultures from adult S100B-EGFP^+^ mice were fixed and stained to characterize neuronal, glial, telocyte-like, and pacemaker cell populations. Antibodies against βIII-tubulin, GFAP, Iba1, c-Kit, and HCN4 were used alongside EGFP^+^ fluorescence. Nuclei were visualized with NucBlue to contrast the endogenous EGFP^+^ signal. Confocal microscopy enabled spatial mapping of cell types and their interactions, as well as measurement of cluster dimensions.

### Flow cytometry

Dissociated sinoatrial node tissue (and right atrium) from S100B-EGFP^+^ mice and non-transgenic C57BL/6J controls was acquired on a BD flow cytometer (FACSDiva 9.0.1); spillover compensation used the acquisition-stored matrix, and gating and quantification were performed in Python (FlowKit). Cells were gated sequentially on forward- and side-scatter, singlets (FSC-H versus FSC-A), and EGFP^+^ fluorescence (FITC channel). The EGFP^+^ threshold was defined as the 99.9th percentile of the paired non-transgenic control distribution and applied identically to reporter and control, because autofluorescence of cardiac tissue in the FITC channel overlaps the dim edge of the EGFP^+^ population; reported EGFP^+^ frequencies are expressed as the excess above this control-defined baseline. Viability was assessed by propidium iodide exclusion, and the EGFP^+^ fraction was quantified as a proportion of singlets.

### Microelectrode array recordings

Electrical activity of SAN-derived cultures was recorded with a high-density CMOS-based MEA system (3Brain CorePlate) with 4,096 electrodes. Because these CMOS arrays are opaque and incompatible with live imaging, MEA recordings were performed on a separate set of cultures grown directly on the array, in parallel with the imaged preparations rather than on the same dishes. Cultures were maintained on extracellular matrix coatings and recorded under physiological conditions without stimulation. Spontaneous spike detection and network synchrony were analyzed.

### SAN tissue explant culture on soft hydrogels

Intact SAN explants from adult S100B-EGFP^+^ mice were micro-dissected and cultured directly on soft hydrogel-coated dishes (~1 kPa) without enzymatic dissociation, in serum-supplemented DMEM/F12 at 37 °C and 5% CO_2_ for up to seven days. Cell outgrowth and tissue viability were monitored by transmitted light and fluorescence microscopy, with careful medium changes to preserve the substrate.

*Detailed step-by-step protocols, including digestion parameters, substrate preparation, and imaging settings, will be published in a peer-reviewed journal*.

## Results

### Dissociated SAN cultures contain a morphologically and behaviorally heterogeneous S100B-EGFP^+^ population

Dissociation of adult mouse SAN produced a heterogeneous primary culture containing NeuN^+^ neuronal-like cells, GFAP^+^/S100B^+^ glia and F4/80^+^ macrophages.

Live imaging identified at least six reproducible morphological and behavioral classes within the S100B-associated cell population (Table 1, Fig. 2). These included elongated, spindle-shaped, directed-motile, amoeboid, meandering, granular engulfing, long-extension network-forming, and stationary perinuclear-bright phenotypes. Representative time-lapse sequences show mitotic division, directed migration, dynamic extension and retraction of long cellular projections, remodeling of intercellular contacts, and engulfment of cellular debris (Fig. 3). In a subset of cells, the EGFP^+^ signal overlapped the nuclear (SPY650-DNA) channel, consistent with either nuclear S100B localization or passive nuclear distribution of free EGFP^+^.

**Table 1.** A morphological and behavioral framework for the S100B-associated cell population.

| Phenotype | Morphological features | S100B-EGFP<sup>+</sup> fluorescence | Behavior in culture | Possible identity |
| --- | --- | --- | --- | --- |
| **Elongated bright cell** | Elongated, length exceeding width, smooth edges, few or no lateral branches. | Bright, uniform cytosolic signal along the long axis, occasionally intensified near the nucleus. | Moderately motile with predominantly directed movement. | Non-myelinating Schwann cell [31,32,33] |
| **Highly plastic spindle cell** | Slender, elongated soma alternating between stretched and spread forms, occasional thin protrusions. | Bright, uniform signal along the long axis, sometimes concentrated near the nucleus during division. | Moderately motile with persistent movement; division observed in uni- and binucleate states. | Proliferating Schwann cell [34,35] |
| **Amoeboid with protrusions** | Slender to rounded, highly plastic body with short arms, occasional ruffles or blebs. | Consistently bright; diffuse, slightly higher along the cell body than extensions. | Highly motile with meandering paths; frequently contacts and partially envelops neighbours. | Macrophage or phagocytic Schwann cell [36,37,45] |
| **Long-extension networker** | Slender soma with one or more elongated, thin extensions contacting distant cells. | Diffuse along soma and extensions, intensified at contact sites; blebs on elongations. | Highly mobile, with extensions that extend, retract, and form intercellular bridges. | Peri-synaptic Schwann cell or fibroblast [38,39] |
| **Stationary perinuclear-bright cell** | Round to oval soma with a dynamic cytoplasmic border around a central nucleus. | Bright perinuclear signal with a diffuse cytoplasmic halo. | Non-migratory over several days, anchored, with slow pulsatile boundary changes. | Satellite glial cell or quiescent Schwann cell [41,42] |
| **Granular engulfing cell** | Small soma with irregular outline and punctate cytoplasmic signal. | Discrete, vesicle-like puncta with increased signal after engulfment. | Highly motile, frequent interactions, evidence of engulfing material from apoptotic cells. | Phagocytic Schwann cell or macrophage [37,46] |

*Six phenotypes distinguished by morphology under live fluorescence microscopy, S100B-EGFP<sup>+</sup> distribution, and live-cell behavior. Bracketed citations in the final column indicate published cell types whose reported morphology and behavior are consistent with each phenotype. Proposed identities are provisional inferences from morphology and behavior and await molecular confirmation. Phenotypes were classified from long-term time-lapse microscopy [40].*

**Fig. 1.**
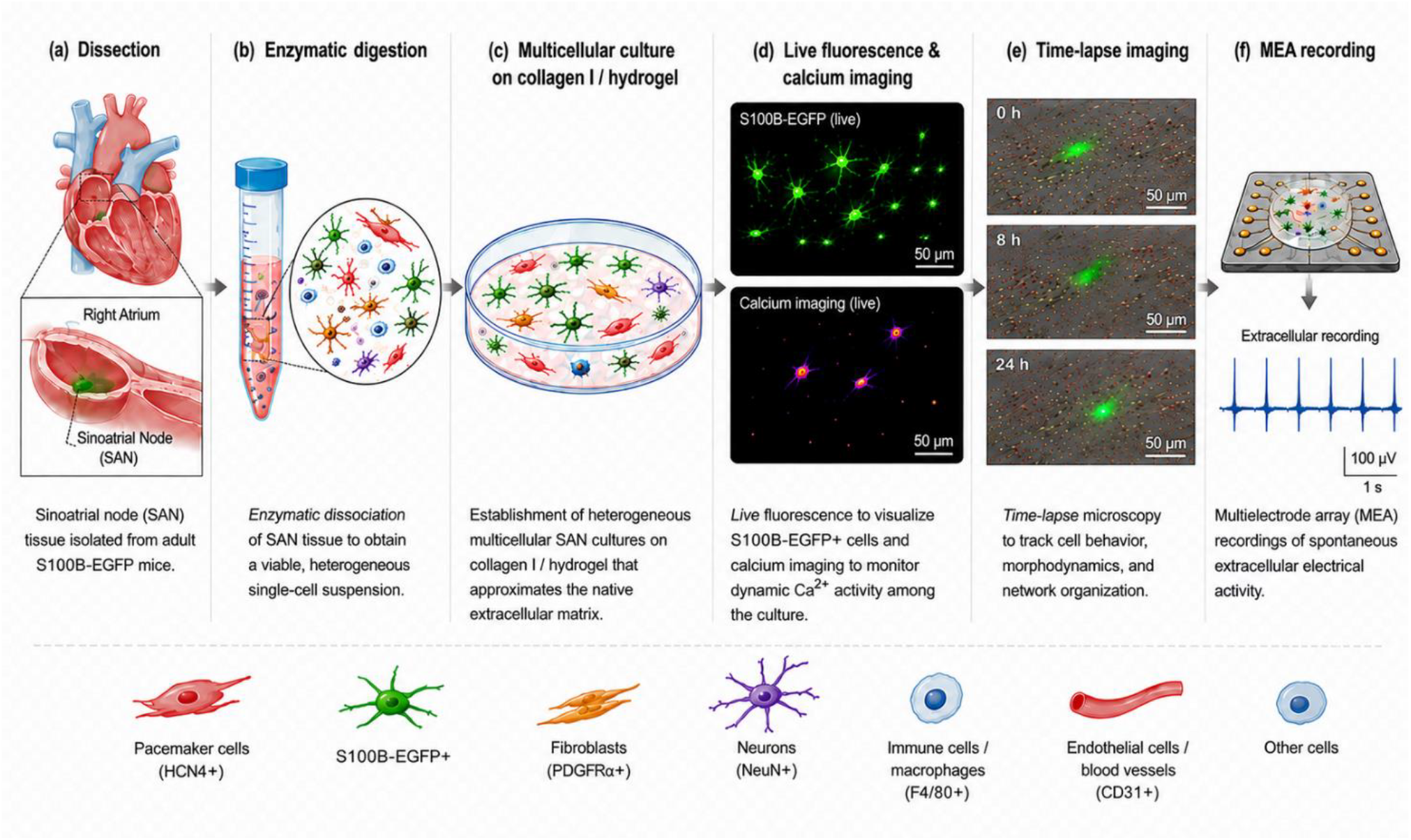
Overview of the model pipeline. **Establishment and characterization of an adult mouse sinoatrial node (SAN) multicellular culture platform. (a)** Microdissection of the SAN from the right atrium of adult S100B-EGFP^+^ mice. **(b)** Enzymatic dissociation of SAN tissue to generate a viable heterogeneous single-cell suspension. **(c)** Establishment of multicellular primary SAN cultures on collagen I hydrogel. **(d)** Live imaging of cultured cells using endogenous S100B-EGFP^+^ fluorescence, and separately, calcium imaging to identify spontaneous calcium activity within the multicellular network. **(e)** Time-lapse microscopy for longitudinal assessment of cell behavior, migration, and network organization. **(f)** Multielectrode array (MEA) recordings with spontaneous extracellular electrical activity in cultured SAN cells.

**Fig. 2.**
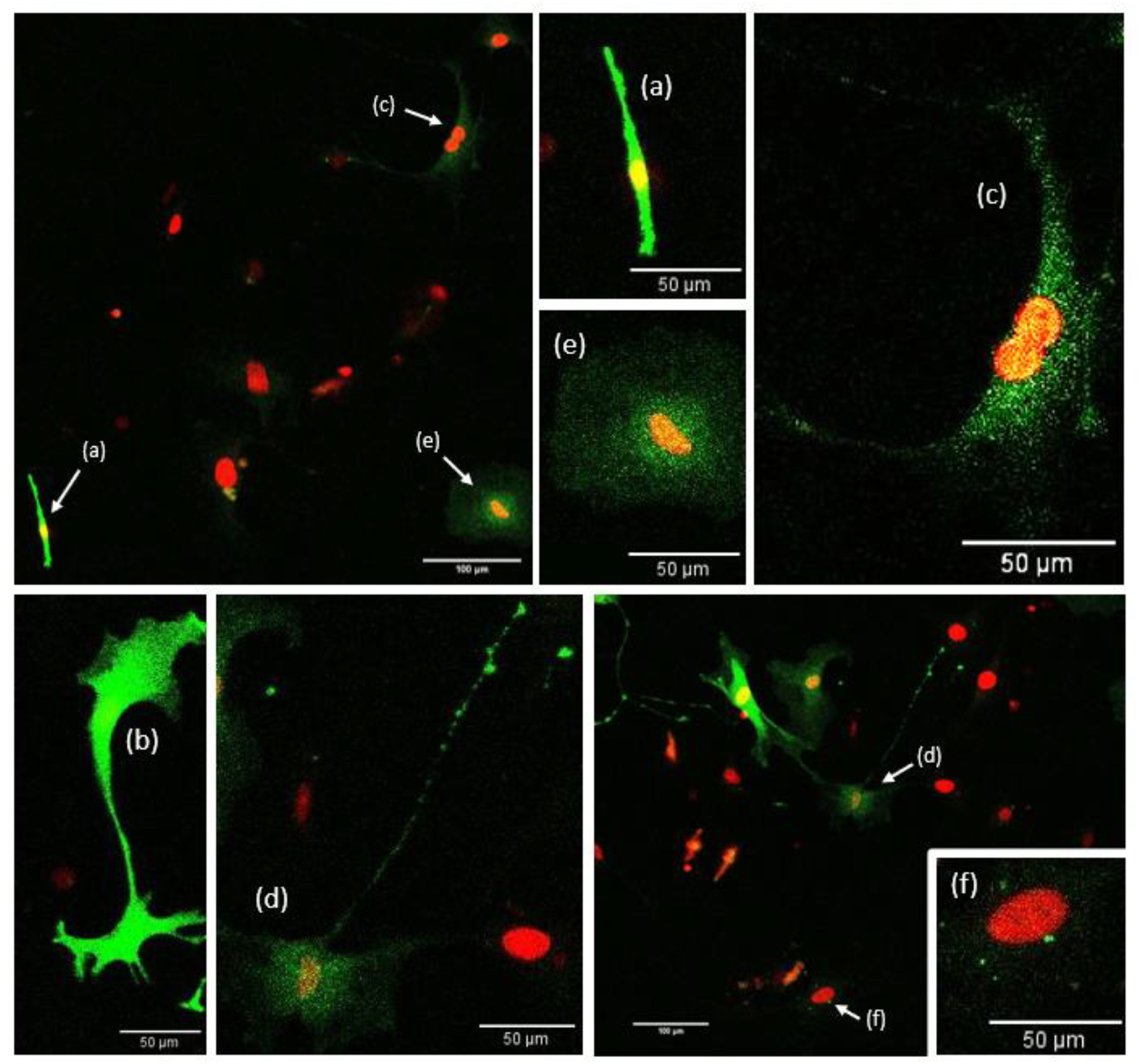
Morphological and fluorescence-intensity heterogeneity of S100B^+^ cells. **Live confocal imaging of S100B-EGFP**^**+**^**-positive cells in multicellular SAN cultures**. Green, endogenous S100B-EGFP^+^; red, nuclei (SPY650-DNA, far-red, pseudocolored red). Subtypes distinguished by morphology and EGFP^+^ intensity: **(a)** elongated bright cell, thin smooth soma with uniform signal along the long axis; **(b)** highly plastic spindle cell with short protrusions; **(c)** amoeboid cell with larger body and diffuse cytosolic signal; **(d)** long-extension networker projecting a thin, beaded process toward distant cells; **(e)** stationary perinuclear-bright cell with bright perinuclear signal and diffuse halo; **(f)** small granular cell with punctate cytosolic signal. Scale bars, [X] µm. Full morphological, intensity, and behavioral criteria in Table 1; dynamic behaviors (migration, division, engulfment) are shown by time-lapse in Figure 3.

**Fig. 3.**
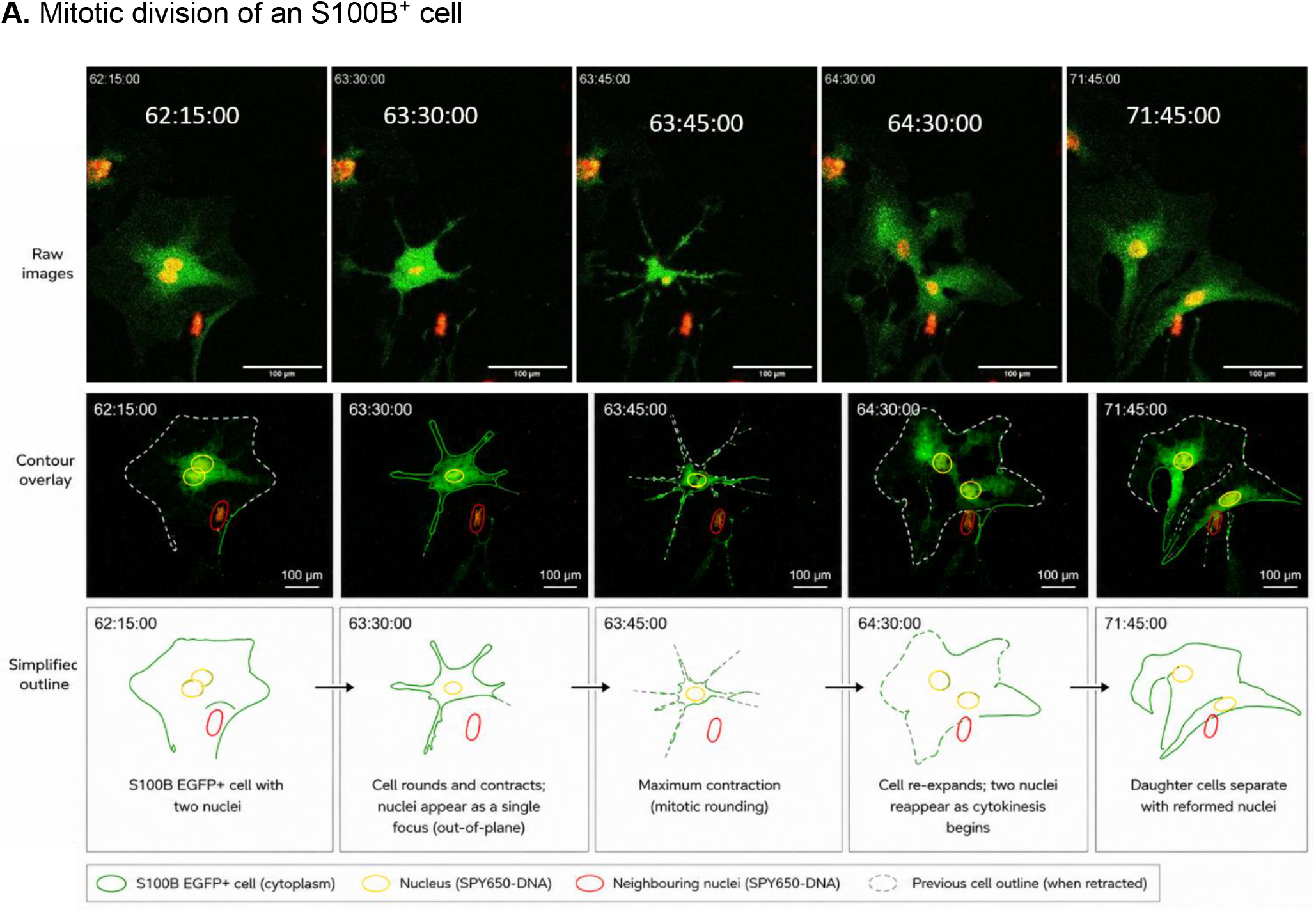
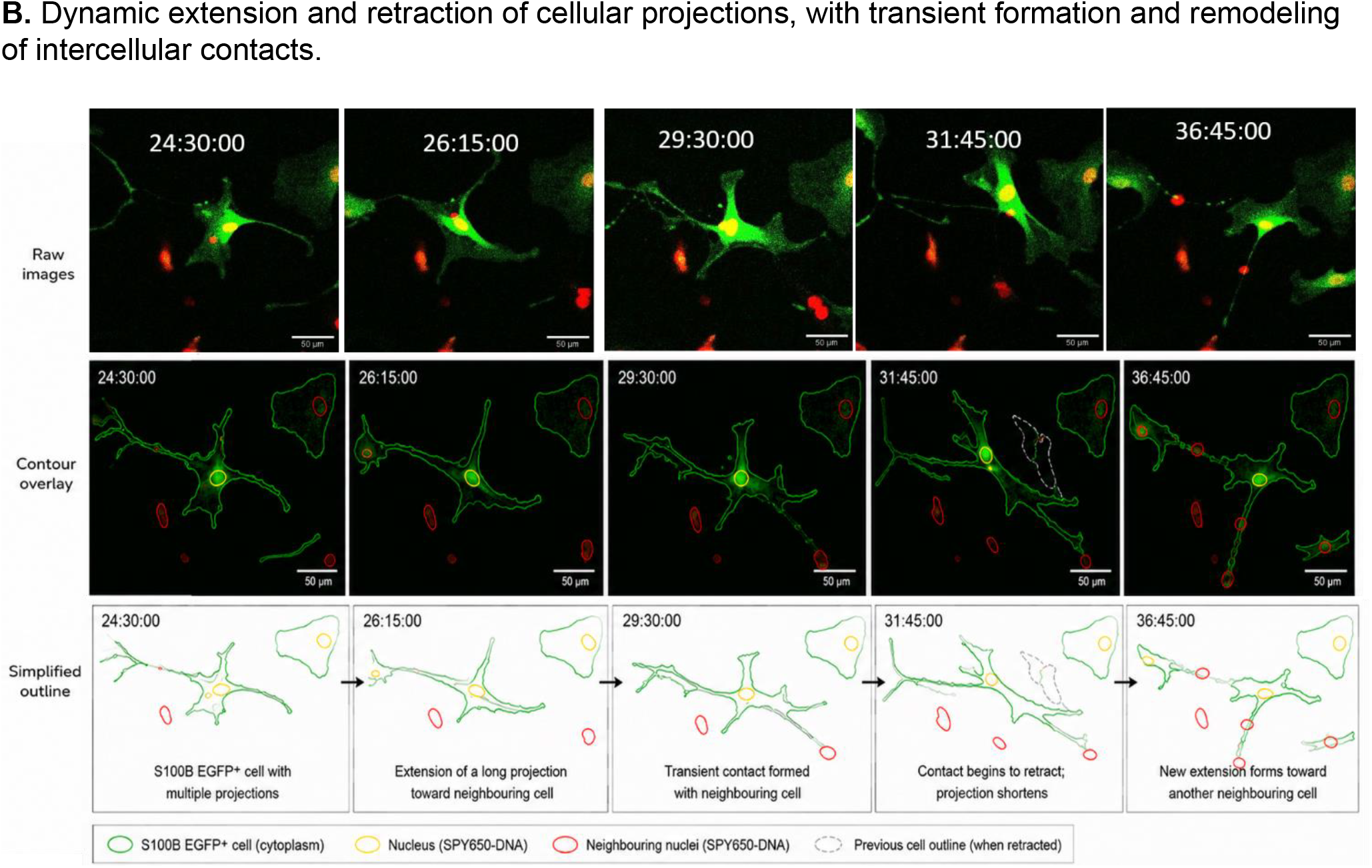
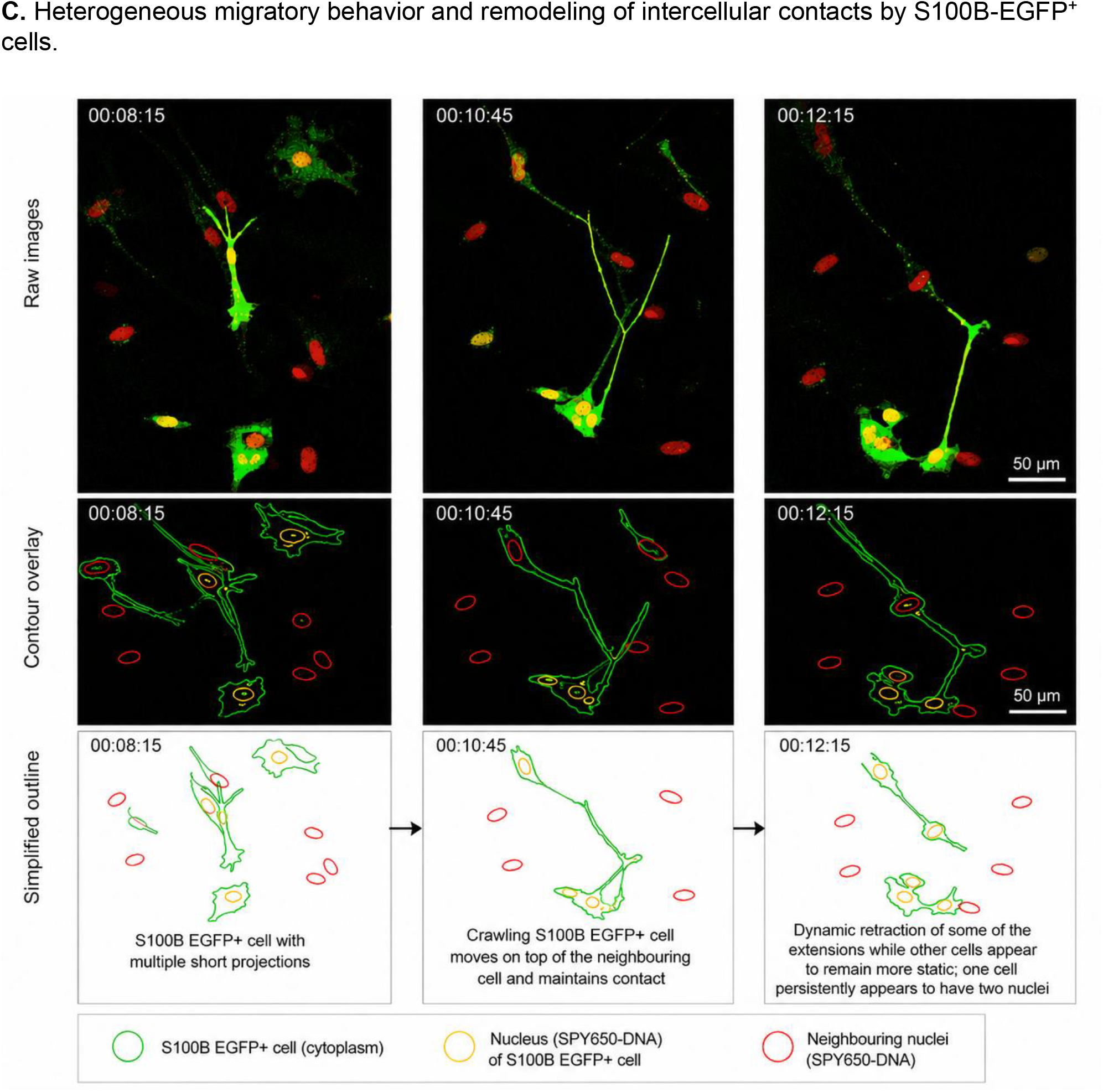
Representative dynamic behaviors of S100B^+^ cells revealed by live imaging. **Representative time-lapse sequences illustrating dynamic behaviors of S100B-EGFP**^**+**^ **cells in multicellular sinoatrial node cultures. (A)** An S100B-EGFP^+^ cell undergoes mitotic division. **(B)** An S100B-EGFP^+^ cell continuously extends and retracts long cellular projections while forming transient contacts with neighboring cells. **(C)** S100B-EGFP^+^ cells exhibit heterogeneous dynamic behaviors within the same field, with some cells remaining relatively stationary while others undergo pronounced migration accompanied by remodeling of cellular processes and intercellular contacts. One S100B-EGFP^+^ cell persistently appears to contain two nuclei throughout the imaging period.

### S100B-EGFP^+^ cells comprise a defined fraction of the dissociated node

To estimate the abundance of S100B-expressing cells in the node, enzymatically digested SAN tissue from S100B-EGFP^+^ mice was analyzed by flow cytometry, with non-transgenic C57BL/6J mice used to set the EGFP^+^ gate. Relative to non-reporter controls, in which EGFP^+^ events were near-absent (≤0.1% of single cells), S100B-EGFP^+^ SAN tissue contained a distinct EGFP^+^ population comprising approximately 9% of single cells (mean 9.1%, range 6.4–10.6%, n = 6; Fig. 4), with a comparable fraction in right atrial tissue (mean 8.8%, range 7.4–10.1%, n = 6; Figs. 4, 5). The EGFP^+^ threshold was defined as the 99.9th percentile of the paired non-transgenic control distribution and applied identically to reporter and control (Methods). Total event yields were low, consistent with the small size of the node.

**Fig. 4.**
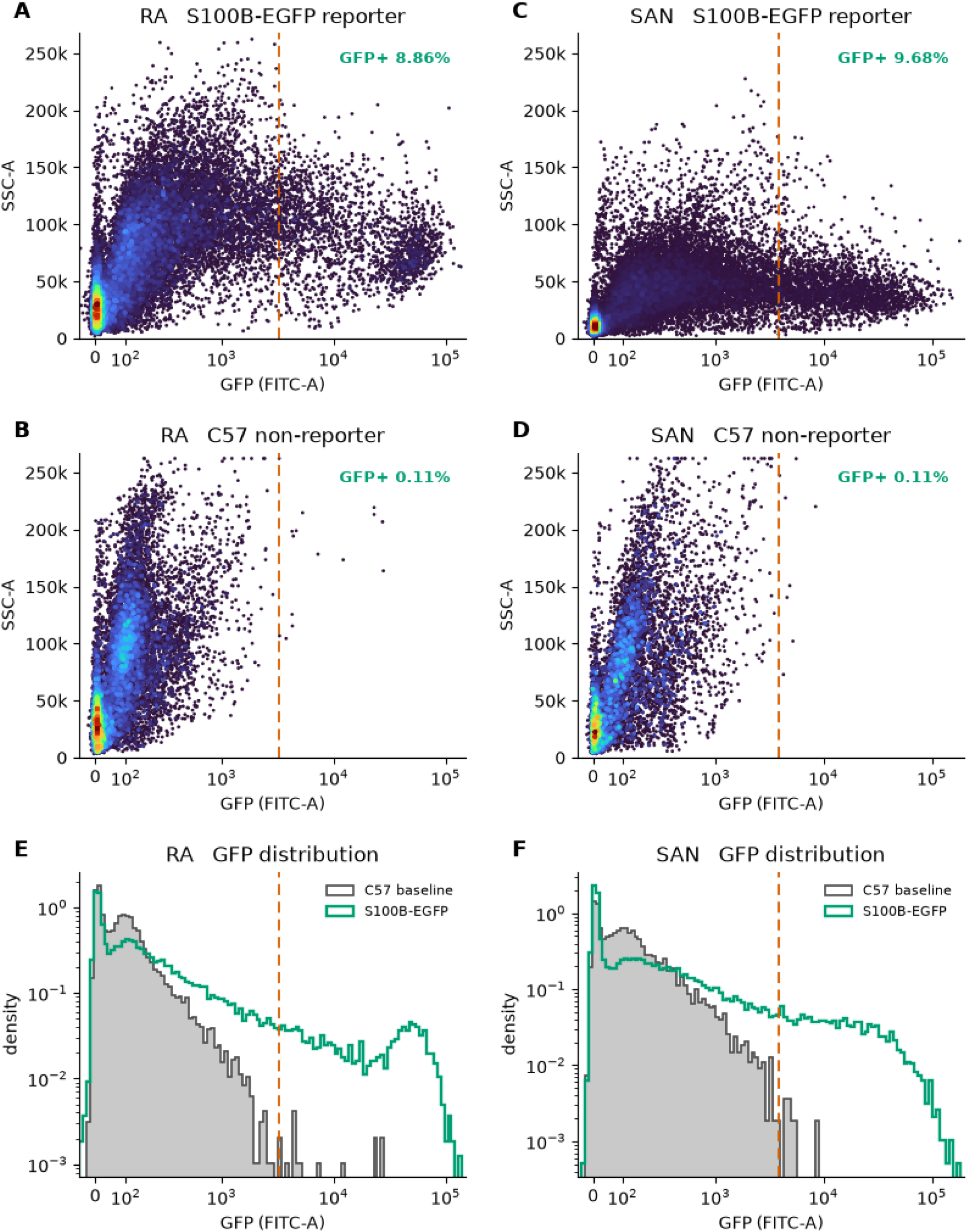
The S100B-EGFP^+^ reporter identifies a discrete EGFP^+^-positive population in dissociated right atrium and sinoatrial node. Flow cytometry of enzymatically dissociated right atrium (RA) and sinoatrial node (SAN) from S100B-EGFP^+^ reporter mice and non-transgenic C57BL/6JN controls, aged 5–6 months. All plots show single, viable (propidium iodide–negative) cells after sequential exclusion of debris (FSC-A/SSC-A), doublets (FSC-A/FSC-H), and dead cells. Spillover compensation was applied using the acquisition-stored matrix, and gating and quantification were performed in Python (FlowKit). (**A–D**) SSC-A versus EGFP^+^ (FITC-A) density plots for reporter (**A**, RA; **C**, SAN) and control (**B**, RA; **D**, SAN). The orange dashed line is the EGFP^+^-positive threshold, defined as the 99.9th percentile of the paired C57BL/6JN control EGFP^+^ (FITC-A) distribution and applied identically to reporter and control; by construction, ~0.1% of control cells fall above it. Values give EGFP^+^ positive cells as a percentage of viable single cells for the representative sample displayed (**A** 8.86%, **C** 9.68%). (**E, F**) EGFP^+^ (FITC-A) density histograms overlaying the C57BL/6JN baseline (grey, filled) and S100B-EGFP^+^ reporter (green) for RA (**E**) and SAN (**F**), with the same control-defined threshold (orange dashed). Reporter samples show a right-shifted EGFP^+^-high population extending to ~10^5^ (FITC-A) that is absent from control. Across biological replicates, EGFP^+^-positive fractions (percentage of viable single cells; per-sample gated events 18,044–40,187) were: RA reporter 8.83% (range 7.39–10.12%, n = 6), SAN reporter 9.09% (range 6.36–10.64%, n = 6). Non-transgenic controls were 0.11% (RA, 13,277 events) and 0.11% (SAN, 7,215 events); a single control baseline was acquired per tissue. Because cardiac tissue autofluoresces in the FITC channel, positivity is defined relative to the non-transgenic control rather than by an absolute gate, and the reporter signal is interpreted as the excess above this baseline.

**Fig. 5.**
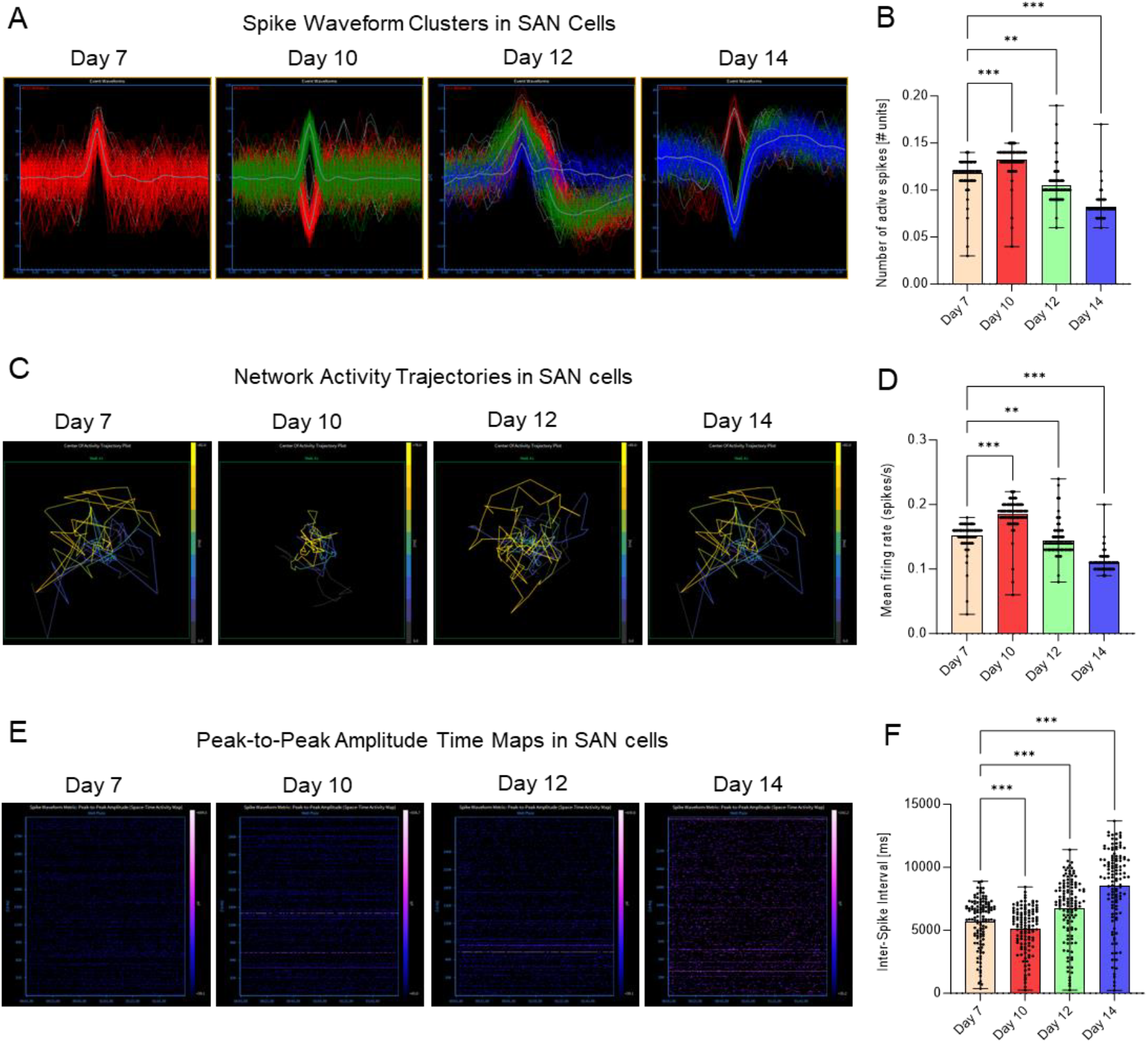
Electrophysiological activity and network dynamics of SAN cultures across days 7–14. **(A)** Representative spike waveform clusters from extracellular recordings; thin lines, individual spikes; bold, cluster mean; polarity differences reflect electrode orientation and coupling. **(B)** Active spikes increase from day 7 to 10 (p < 0.001), then stabilize or decline at days 12 to 14, indicating peak excitability at day 10. **(C)** Center-of-activity trajectories: sparse at day 7, coherent at day 10, dispersed at days 12 to 14. **(D)** Mean firing rate (spikes/s): maximal at day 10 versus day 7 (p < 0.001); reduced at days 12 and 14 (p < 0.01 to p < 0.001 versus day 10).**(E)** Peak-to-peak amplitude maps: sparse and localized at day 7; widespread and synchronized at day 10; reduced at days 12 to 14. **(F)** Inter-spike interval: shorter at day 10 versus day 7 (p < 0.001); longer at days 12 and 14 versus day 10 (p < 0.001). Data are mean ± SEM; one-way ANOVA with Bonferroni post hoc.

### Multicellular SAN cultures remain spontaneously electrically active

Ca^2+^ imaging with Calbryte 520 revealed spike-like Ca^2+^ transients consistent with action-potential-coupled events by day 5 in vitro, providing an optical surrogate for electrical activity in these imageable cultures, which cannot be recorded on the opaque MEA arrays.

In separate experiments, extracellular MEA recordings on days 7, 10, 12, and 14 showed multi-unit spiking with stable, polarity-diverse waveform clusters (Fig. 5). Network coherence and firing rate peaked on day 10, when center-of-activity trajectories were most organized and inter-spike intervals were shortest. Activity persisted on days 12 to 14, with reduced amplitude and synchrony. Peak-to-peak amplitude maps and trajectory videos (Supplementary Videos) show sparse activity on day 7, widespread synchronized bursts on day 10, and partial desynchronization thereafter. Spontaneous electrical activity, the defining functional property of nodal tissue, persisted beyond ten days without external stimulation.

### Spontaneous formation of S100B^+^-enriched three-dimensional clusters

Primary SAN cultures spontaneously formed three-dimensional, roughly spherical multicellular clusters without external patterning cues or scaffolding, typically within three to five days in vitro. Immunofluorescence showed that these clusters contained both HCN4^+^ pacemaker-associated cells and S100B-EGFP^+^ cells, with S100B-EGFP^+^ cells present at both the core and periphery (Supplementary Fig. S1). Some clusters exhibited spontaneous contractile activity, although contraction was not causally linked to the S100B-EGFP^+^ population in these experiments. Clusters were absent immediately after seeding and emerged progressively, indicating self-organization rather than pre-formed aggregation. They grew to confocal-measured thicknesses of approximately 17 to 32 µm (Supplementary Fig. S2). The integration of S100B^+^ cells within these organized structures points to a potential role for non-myocyte populations in supporting multicellular organization in SAN-derived cultures; S100B-expressing cells likewise self-organize into parenchymal clusters in other tissues [30].

## Discussion

Rather than reducing the SAN to isolated cell populations, this model allows direct observation of cellular interactions within an intact, spontaneously active multicellular preparation. We established a multicellular culture model of the adult sinoatrial node that remains viable and electrically active outside the body for more than ten days. Adult pacemaker myocytes survive only briefly once isolated, remaining viable for approximately 24–48 hours in culture [43], and conventional approaches recover individual cell populations at the expense of the multicellular context that shapes nodal function. In contrast, our cultures retain a cellular environment containing HCN4^+^ cells together with fibroblasts, neurons, glia, immune cells, and S100B-associated cells, while maintaining spontaneous electrical activity for more than ten days. The compliant collagen–gelatin hydrogel, tuned to approximate the mechanical properties of the native SAN matrix, was central to maintaining this activity without lineage-specific media or exogenous enrichment. This preserved multicellular context allows rare S100B-associated cells to be followed over days within an intact, spontaneously active network, rather than studied in isolation.

Within this system, the S100B-associated cell population exhibits marked morphological and behavioral heterogeneity, as evidenced by live time-lapse fluorescence imaging. Several phenotypes resemble reported peripheral glial morphologies, including elongated migratory and network-forming cells, whereas others display amoeboid or phagocytic behaviors reminiscent of macrophage-like states. These observations are based on morphology and behavior and do not establish lineage identity but instead motivate future characterization of this population. Formal identification of these cells will require complementary molecular approaches, in particular single-cell RNA sequencing, which this imaging-based platform does not provide. The ongoing debate over whether resident macrophages contribute to nodal conduction [10,11] illustrates the difficulty of assigning functional roles to morphologically defined cell populations *in situ*. The platform supports repeated time-lapse imaging of the same cultures over days, allowing these hypotheses to be tested directly by tracking and perturbing individual cells over time.

Another notable feature was spontaneous self-organization. Without external patterning, a subset of cells assembled into three-dimensional clusters containing both HCN4^+^ and S100B-EGFP^+^ cells, whereas others remained dispersed. Some clusters also exhibited spontaneous contractile activity. Their cellular composition suggests that the model retains aspects of the native SAN microenvironment. This is relevant because the native SAN is itself a three-dimensional meshwork in which pacemaker and non-myocyte populations are interwoven. Whether S100B-associated cells actively scaffold these structures or are passively incorporated remains unresolved, but the platform provides a tractable system for investigating how this architecture emerges and how non-myocyte populations contribute to its assembly.

By bridging intact SAN preparations and isolated-cell cultures, this model offers a practical system for the longitudinal study of rare S100B-associated cells within a spontaneously active multicellular environment, and for resolving how non-myocyte populations shape sinoatrial node function and its decline with age.

## Limitations

Several limitations stem from this study’s design. First, dissociation and culture induce phenotypic changes, and we characterize the model functionally and morphologically rather than asserting molecular equivalence to native tissue. Second, the cultures are mixed populations without lineage tracing, so subtype identities in Table 1 are inferred from morphology, behavior, and reporter intensity and remain provisional. Third, MEA recordings capture network-level extracellular activity and, on their own, do not attribute firing to a specific cell type; the relationship between S100B^+^ cells and the recorded activity is correlative at this stage. Fourth, the explant outgrowth observations are descriptive. Finally, all data derive from adult mouse SAN tissue; extrapolation to human tissue remains to be established.

## Acknowledgements

This research was supported [in part] by the Intramural Research Program of the National Institutes of Health (NIH). The contributions of the NIH author(s) are considered Works of the United States Government. The findings and conclusions presented in this paper are those of the author(s) and do not necessarily reflect the views of the NIH or the U.S. Department of Health and Human Services. We thank Khalid Chakir for breeding the S100B-EGFP^+^ (kosmos) reporter mice, and Rostislav Bychkov for his guidance in sinoatrial node preparation and calcium imaging and for his early studies of S100B in intact SAN tissue.

## Author contributions

G.L.B. conceived the study, designed the culture platform, coordinated the collaboration, performed and analyzed experiments, and wrote the manuscript. R.E.M. contributed to the study concept and to tissue dissociation and primary culture. B.D.Z. refined and optimized the enzymatic dissociation of the small sinoatrial node through successive iterations and supported primary culture. S.M.T. R. performed live-cell confocal imaging, image analysis and helped refine the study concept. P.P. performed immunostaining and confocal imaging and helped refine the study concept. D.B. performed immunostaining and advised on pacemaker myocyte preservation in culture. R.T. contributed to live-cell imaging and to immunostaining of fixed preparations. C.D. performed flow cytometry. D.T. performed preliminary flow cytometry and exploratory sequencing experiments. S.A. set up the microelectrode array recordings and analyzed initial electrophysiological data. K.-W.G.L. performed transcriptomic experiments and analyzed and interpreted the data. A.-M.Z. advised on glial and neuronal biology and established neuronal culture conditions. P.K., M.-H.S., N.H.G., S.D., P.S., and A.B.H. provided reagents, expertise, and analytical resources and contributed to interpretation. R.d.C. provided supervision and resources. E.G.L. supervised the study, provided resources and funding, and contributed to the study concept. All authors reviewed and approved the manuscript.

## Competing interests

The authors declare no competing interests.

## Data availability

Data supporting the findings of this study are available from the corresponding author on reasonable request. Detailed step-by-step protocols will be made available on peer-reviewed publication.

## Notes

### Competing Interest Statement

The authors have declared no competing interest.

